# Apical extrusion prevents apoptosis from activating an acute inflammatory program in epithelia

**DOI:** 10.1101/2023.02.12.528237

**Authors:** Kinga Duszyc, Jessica B. von Pein, Divya Ramnath, Denni Currin-Ross, Suzie Verma, Fayth Lim, Matthew J. Sweet, Kate Schroder, Alpha S. Yap

## Abstract

Apoptosis is traditionally considered to be an immunologically silent form of cell death. Multiple mechanisms exist to ensure that apoptosis does not stimulate the immune system to cause inflammation or autoimmunity. Against this expectation, we now report epithelia are programmed to provoke, rather than suppress, inflammation in response to apoptosis. We found that an acute inflammatory response led by neutrophils occurs in zebrafish and cell culture when apoptotic epithelial cells cannot be expelled from the monolayer by apical extrusion. This reflects an intrinsic circuit where ATP released from apoptotic cells stimulates epithelial cells in the immediate vicinity to produce IL-8. As the epithelial barrier is compromised when apical extrusion fails, this juxta-apoptotic proinflammatory pathway may represent an early-response mechanism at sites of potential microbial ingress. Conversely, apical extrusion prevents inappropriate epithelial inflammation by physically eliminating apoptotic cells before they can activate this proinflammatory circuit.

## Introduction

Dying cells have a potent capacity to provoke inflammation and autoimmune responses if their internal contents are exposed to the extracellular space^1,2^. Indeed, lytic forms of cell death, such as pyroptosis, are characteristically inflammatory^3–5^. In contrast, apoptosis is traditionally considered to be immunologically silent^6,7^, a feature that is important given its ubiquitous presence in healthy tissue turnover. Developmentally regulated apoptosis participates in shaping organs such as the digits of the limb^8–10^, while it has been estimated that 0.5% of the body’s cells undergo apoptosis on daily basis in post-embryonic life^11^. This immunological silence is enforced by diverse mechanisms that provide multiple layers of protection when apoptosis occurs. A key mechanism is the cell-intrinsic process of apoptotic fragmentation, where cytosolic contents are packaged into membrane-bound apoptotic bodies^6,12^, which can then be cleared by professional phagocytes, such as macrophages, or non-professional phagocytes, such as epithelial cells^6,13–19^. In addition, ingestion of apoptotic cells directs professional phagocytes towards an anti-inflammatory and immunosuppressive phenotype, secreting anti-inflammatory cytokines, such as TGF-β1^20,21^, and downregulating pro-inflammatory mediators^21,22^. Together, these defences ensure that apoptosis is not only immunologically silent, but also an actively anti-inflammatory type of cell death.

Epithelia have an additional line of response to potentially shield the immune system from apoptotic cells. This is the morphogenetic process of apical extrusion, where sporadic apoptotic cells are physically expelled by their neighbours into the external environment^23–30^. Live cell imaging in cell culture and zebrafish embryos indicates that this is a rapid process, with apoptotic cells often completing their fragmentation process only after they have been expelled^26^. Apical extrusion is intrinsic to the epithelium itself, as it can be executed by epithelial mono-cultures, and also seals the epithelium as it expels the apoptotic cells^23,26^, further separating them from immune detection. Since epithelial barriers are subject to diverse environmental stresses that will injure cells, apical extrusion is well-placed to serve as a first responder to apoptosis in these tissues.

Despite these attractive design features, we do not yet know whether there are inflammatory consequences if apical extrusion fails. Perhaps other homeostatic processes, such as efferocytosis, compensate to preserve the immunological invisibility of apoptotic cells. In this study, we address this question in zebrafish embryos and cell culture models. We report that, against the traditional view, epithelia activate an acute inflammatory response when apoptotic cells cannot be eliminated by apical extrusion. This response reflects an intrinsic purinergic cell-cell signaling pathway in epithelia, where apoptotic cells stimulate their neighbours to produce IL-8. As extrusion failure disrupts the epithelial barrier, this may represent an early-warning defence that is engaged at sites of potential microbial ingress. Conversely, apical extrusion prevents sporadic apoptosis from provoking inflammation inappropriately by physically eliminating apoptotic cells before they can activate this signaling pathway.

## Results

### Apoptotic epithelial cells stimulate neutrophil recruitment when they cannot be eliminated by apical extrusion

We used zebrafish embryos as a vertebrate model to assess the inflammatory impact of epithelial apoptosis. Sporadic apoptosis was induced by laser-microirradiating small clusters of cells (≤ 4 cells) in the 4 days post-fertilization (dpf) zebrafish periderm, the outermost epidermal layer of the embryo. We measured the recruitment of neutrophils to gauge the early inflammatory response, as these are one of the first cellular responders to perturbations of homeostasis. To identify neutrophils at the periderm, heterozygous *Tg(mpx:mCherry*)^uqay3^ carriers, which express mCherry in neutrophils, were crossed with *Tg(krt4:GFP-CAAX)*^uqay1^ animals, expressing membrane-targeted GFP in the periderm, and double positive offspring were selected for experiments (Fig 1A). As recently reported^26^, microirradiated peridermal cells underwent apoptosis, identified by cell rounding and blebbing. Within 10-15 min whole apoptotic cells were eliminated from the periderm into the external environment by apical extrusion. This was associated with rearrangement of the surrounding neighbour cells to form a rosette sealing the epithelium at the site of apoptotic extrusion. Induction of apoptosis in these animals was not accompanied by an acute inflammatory response, as scant neutrophils were evident near the sites of apoptosis (Fig 1B,C, Video S1). Therefore, epithelial apoptosis was immunologically silent under these conditions, where the periderm was able to mount an effective extrusion response.

**Fig 1.**
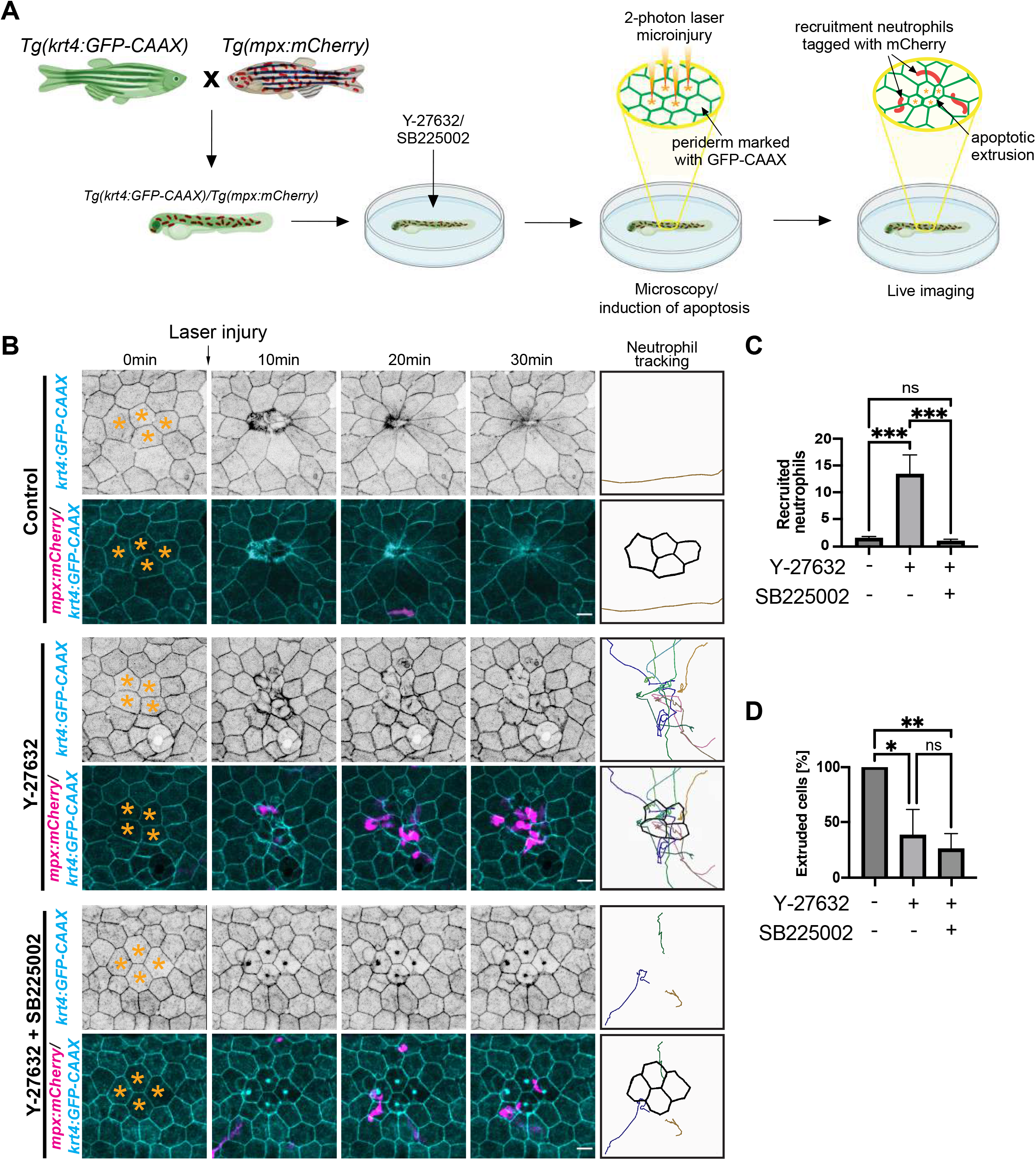
Extrusion failure causes apoptotic epithelial cells to recruit neutrophils. **A**. Schematic diagram of zebrafish mating and experimental set up to induce peridermal apoptosis and live image apoptotic extrusion and neutrophil recruitment. **B**. Selected frames from live-imaging of apoptosis and neutrophil recruitment to the sites of apoptosis in 4dpf zebrafish periderm. Prior to imaging zebrafish were treated with 45μM Y-27632 and 100nM SB225002 (16hrs). Yellow asterisks – apoptotic cells. Panels on the right represent tracks of moving neutrophils within the imaging fields. **C**. Quantification of neutrophils recruited to the sites of apoptosis within 30 min following laser microinjury of 4dpf zebrafish peridermal cells. Prior to imaging zebrafish were treated with 45μM Y-27632 and 100nM SB225002 (16hrs). **D**. Percentage of apoptotic cells extruded after Y-27632 and SB225002 treatment (45μM and 100nM 16hrs treatment respectively) in zebrafish 4dpf. Apoptosis was induced by 2-photon mediated laser microinjury. Scale bars: 15μm. XY panels are maximum projection views from z-stacks. All data are means ± SEM; ns, not significant; *p <0.05, **p < 0.01,***p < 0.001; calculated from n≥3 independent experiments analysed with one-way ANOVA Dunnett’s multiple comparisons test.

Then, we asked how the inflammatory silence of apoptosis was affected when apical extrusion was blocked. We used the Rho kinase inhibitor Y-27632 (45μM) to disrupt the contractile response that is necessary to drive apoptotic extrusion^23,26^, its efficacy in the periderm confirmed by reduction in Myosin IIA at cell-cell junctions (Fig S1A,B). Consistent with studies in cell culture^26^, laser-microirradiated, apoptotic cells were retained in the periderm upon Y-27632 treatment (Fig 1D), confirming that apical extrusion was blocked. Strikingly, this was accompanied by the rapid (10-20 min) and robust recruitment of neutrophils into the periderm (Fig 1B,C, Video S1). By implication, these apoptotic epithelial cells quickly became visible to the innate immune system if they were not eliminated by apical extrusion.

The apical extrusion response serves to preserve the epithelial barrier, as well as physically eliminate apoptotic cells. Therefore, it was possible that neutrophils were being recruited in response to ingress of noxious environmental factors when the barrier was disrupted by extrusion failure. Accordingly, we developed a cell culture system to further test the pro-inflammatory impact of epithelial apoptosis when extrusion was compromised, reasoning that the sterile conditions of tissue culture would eliminate contributions of barrier dysfunction, especially microbial ingress. For this, we used Caco-2 colorectal epithelial cells grown to confluence in pure culture, which allowed us to isolate effects that were intrinsic to the epithelial cells. We postulated that if the epithelium that retained apoptotic cells recruited neutrophils, this would be through a soluble chemokine. So, we collected conditioned media from Caco-2 cultures and tested their ability to recruit isolated human neutrophils in Transwell^®^ migration assays (Fig 2A). As serum is known to induce neutrophil migration, we performed these assays with low FBS (0.5%) concentrations. However, under these conditions we found that etoposide was ineffective for inducing cell death, presumably because its mode of action requires serum-stimulated DNA replication in cells. Accordingly, we used 2mM H_2_O_2_, which damages DNA, to induce apoptosis. H_2_O_2_ efficiently induced apoptosis in cells incubated with media containing either 10% or 0.5% FBS (Fig S2A). Finally, we inhibited apical extrusion with the alternative approach of depleting E-cadherin by RNAi (knock-down, KD), providing a complementary test of the role of apical extrusion (Fig 2B,C; Fig S2B,C).

**Fig 2.**
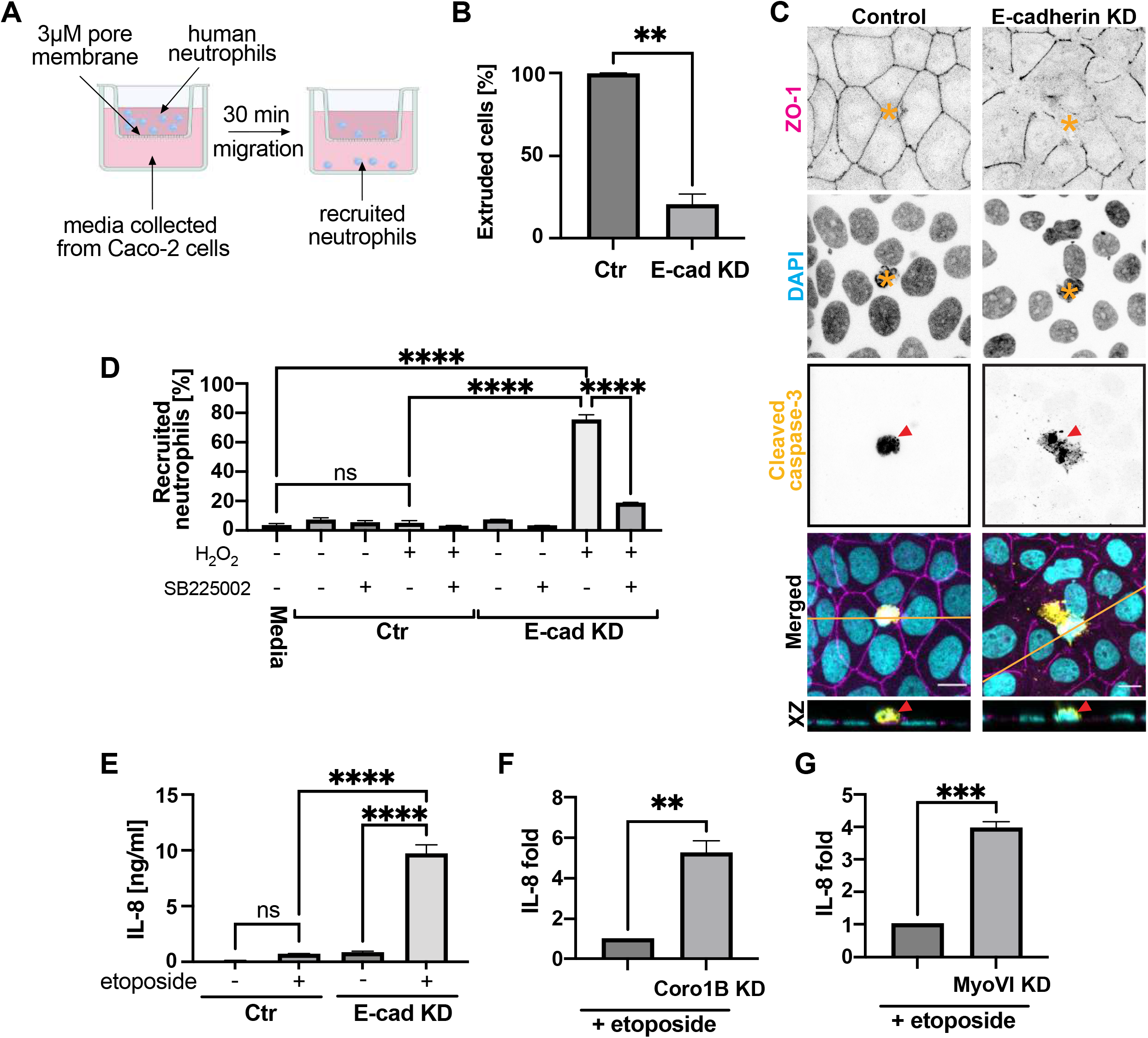
Retained apoptotic epithelial cells recruit neutrophils through IL-8. **A**. Schematic diagram of neutrophil Transwell^®^ migration assay. **B-C**. Effect of E-cadherin knock-down (E-cad KD) on apoptotic extrusion. (B) – quantification; (C) – representative immunofluorescence images (yellow asterisks – location of apoptotic cells; red arrowhead – positive cleaved caspase-3 staining (apoptotic marker); yellow line – location of the XZ views). **D.** Percentage of recruited neutrophils in the Transwell^®^ migration assay (A). Conditioned media were obtained by treating (24hrs) control and E-cad KD Caco-2 cells with 2mM H_2_O_2_ or/and 100nM SB225002 (24hrs). **E**. ELISA-based measurement of IL-8 protein level in media collected from Caco-2 cells after 24hrs treatment with 250μM etoposide. **F-G**. Fold increase in IL-8 production by (F) Coronin1B knock-down and (G) Myosin VI knockdown upon etoposide treatment (250μM, 24hrs). Values were normalised to controls. Scale bars: 15μm. XY panels are maximum projection views from z-stacks. All data are means ± SEM; ns, not significant; **p < 0.01,***p < 0.001, ****p < 0.0001; calculated from n≥3 independent experiments analysed with Student’s t-test (B, F, G) or one-way ANOVA Dunnett’s multiple comparisons test (D, E).

Conditioned media from cultures not exposed to H_2_O_2_ did not stimulate neutrophil migration above the background levels seen with unconditioned media. Nor was neutrophil migration altered by media from H_2_O_2_-treated cultures that retained E-cadherin and were able to execute apical extrusion. However, neutrophil migration was robustly stimulated by media when apoptosis was induced with H_2_O_2_ and extrusion was blocked by E-cadherin KD. This effect was not due to E-cadherin depletion, as neutrophil migration was not stimulated by media from E-cadherin KD cultures that had not been treated with H_2_O_2_ (Fig 2D). Taken with our observations in zebrafish embryos, this suggested that epithelia respond to apoptosis by stimulating, rather than suppressing, an innate immune response.

### IL-8 is secreted by epithelia to recruit neutrophils when apoptotic extrusion is inhibited

What is the chemokine that stimulates migration of neutrophils when extrusion fails? We pursued this question by focusing on CXCL8/IL-8, a classical neutrophil chemokine. First, we used SB225002 to antagonize CXCR2, a major receptor for CXCL8/IL-8. For zebrafish experiments, 3 dpf embryos were pre-treated with 100nM SB225002 as well as Y-27632 (45 μM) for 16 hours before laser microirradiation and imaging at 4 dpf. SB225002 treatment resulted in significantly fewer neutrophils recruited to the sites of failed extrusion (Fig 1B-D, Video S1), demonstrating that CXCR2 was required to stimulate neutrophil migration to epithelia that retained apoptotic cells. This was confirmed in our cell culture system, as SB225002 blocked neutrophil recruitment to conditioned media (Fig 2D). Therefore, CXCR2 was necessary to allow neutrophils to detect epithelial apoptosis when extrusion was blocked.

CXCL8/IL-8 is the most potent ligand of CXCR2, but this receptor is also engaged by CXCL1-3 and CXCL5-8^31,32^. Therefore, we directly tested whether IL-8 production was increased in media from epithelia that retained apoptotic cells. As we were not measuring neutrophil recruitment, we used etoposide to induce apoptosis. There was no significant change in IL-8 production over background when cultures were treated with either etoposide or E-cadherin KD alone. However, IL-8 production was significantly increased in media from E-cadherin KD cultures treated with etoposide (Fig 2E). In contrast, we did not detect production of other common cytokines IL-6 (Fig S2D) or IL-1β (Fig S2E) in these experiments. Importantly, IL-8 production by etoposide-treated cultures was also significantly increased when we blocked extrusion with either Coronin 1B^25^ (Fig 2F) or Myosin VI^26^ (Fig 2G) RNAi, targeting two cytoskeletal proteins that independently support apical extrusion. Therefore, IL-8 is released from epithelia to recruit neutrophils when apoptotic cells cannot be eliminated by apical extrusion.

### IL-8 is produced by the neighbours of retained apoptotic cells

These findings suggested that epithelia possess an intrinsic mechanism that provokes acute inflammation in response to apoptosis. We then asked how apoptotic cells, which engage multiple anti-inflammatory mechanisms^20–22^, instead upregulate the release of pro-inflammatory IL-8 when retained in the epithelial barrier. IL-8 mRNA levels in the monolayers were increased in response to apoptosis to a greater extent in E-cadherin KD cultures than in controls (Fig 3A), implying that its synthesis was transcriptionally upregulated. Furthermore, the release of IL-8 from monolayers was blocked when we inhibited membrane transport with Brefeldin A (Fig S3A). Therefore, the increased release of IL-8 was due to upregulated synthesis and secretion from the monolayers. We then immunostained for IL-8 to identify the cells within the monolayer that were responsible for its production, taking advantage of Brefeldin A to enhance its signal within the cells. Apoptotic cells (detected with cleaved caspase-3) did not show IL-8 staining, nor was it evident in control monolayers that were able to extrude apoptotic cells. Surprisingly, cytoplasmic IL-8 was instead evident in the neighbours of retained apoptotic cells in E-cadherin KD monolayers, typically within 1-2 cell diameters of the apoptotic cells (Fig 3B,C). This implied that the epithelium immediately surrounding the apoptotic cells was the source of IL-8 when extrusion was blocked by E-cadherin KD. We then pursued the pathway that was responsible for upregulating IL-8 production.

**Fig 3.**
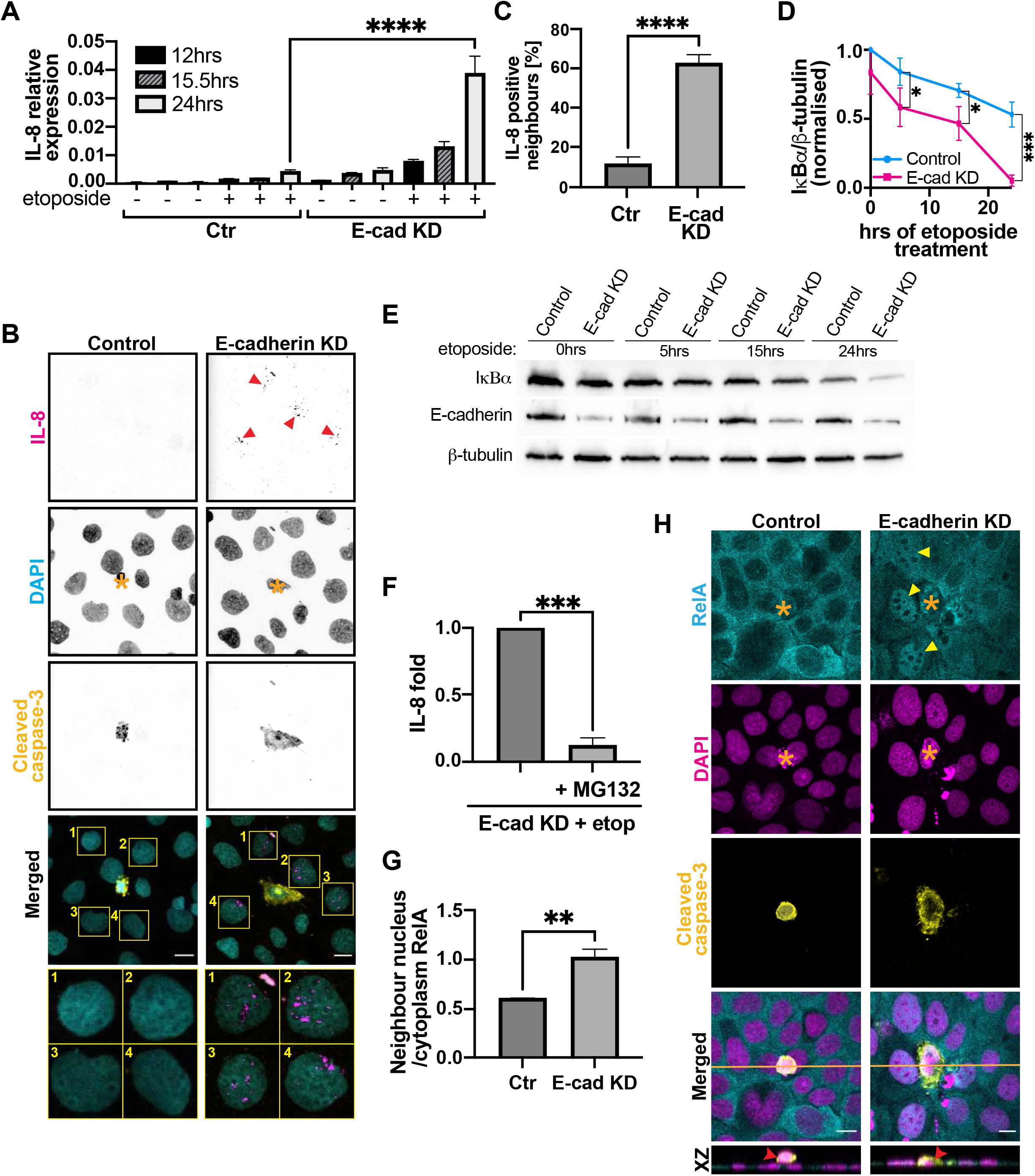
Cells neighbouring retained apoptotic cells are the source of IL-8. **A-B**. Representative immunofluorscent images (A) and quantification (B) of IL-8 positive neighbouring cells upon successful (control cells) and failed (E-cadherin KD cells) apoptotic extrusion. (A) red arrowheads – positive IL-8 staining in cells directly neighbouring the retained apoptotic cell; yellow asterisks – location of apoptotic cells. **C**. IL-8 production (measured by qPCR) in control and E-cadherin KD cells in response to etoposide treatment (250μM); levels of IL-8 mRNA were normalised to the levels of the houskeeping HPRT. **D-E**. Quantification (D) and a representative immunoblot (E) showing degradation of IκBα in control and E-cadherin KD cells upon treatment with etoposide (250μM). **F**. Change in IL-8 production (fold, normalised to E-cadherin KD) upon treatment with proteasome inhibitor MG132 (50μM, 24hrs). Cells in both conditions were treated with etoposide (250μM, 24hrs). **G-H**. Quantification (G) and representative immunofluorescent images (H) of nuclear translocation of RelA (p65 subunit of NF-κB). (H) yellow asterisks – location of apoptotic cells; yellow arrowheads – nuclear RelA staining in cells directly neighbouring the retained apoptotic cell; red arrowheads – location of apoptotic cells; yellow line - location of the XZ views. Scale bars: 15μm. XY panels are maximum projection views from z-stacks. All data are means ± SEM; *p < 0.05, **p < 0.01,***p < 0.001, ****p < 0.0001; calculated from n≥3 independent experiments analysed with Student’s t-test (B, F, G), one-way ANOVA Dunnett’s multiple comparisons test (C) or two-way ANOVA Sidak’s multiple comparisons test (D).

NF-κB is a classic transcriptional activator of IL-8 production that translocates to the nucleus when its inhibitor IκBα undergoes proteasomal degradation in stimulated cells^33–37^. Upon treatment with etoposide, IκBα was degraded more robustly in E-cadherin KD cells compared to the control cells (Fig 3D,E), suggesting that NF-κB might be activated to induce IL-8 production in E-cadherin KD cells. Consistent with this, inhibiting the proteasomal degradation of IκBα with MG132 prevented etoposide from upregulating IL-8 production in E-cadherin KD cells (Fig 3F). To identify the cells with active NF-κB, we then evaluated its nuclear translocation by immunostaining for its RelA/p65 subunit. We did not detect nuclear RelA in the apoptotic cells. Instead, nuclear RelA was evident in the immediate neighbours of the apoptotic cells retained in E-cadherin KD monolayers. In contrast, the closest neighbours of extruded apoptotic cells in control cultures showed cytoplasmic localization of RelA (Fig 3G,H). This observation suggested that NF-κB signaling was enhanced in the neighbours of apoptotic cells that were retained when extrusion was disabled. Taken with pattern of IL-8 immunostaining, these data implied that an NF-κB signaling pathway was being activated in the neighbours of apoptotic cells to generate an epithelial IL-8 signal.

### Enhanced IL-8 production is not due to direct genotoxic damage in neighbour cells

Agents such as etoposide cause apoptosis by inducing DNA damage, which is often broadly-evident within epithelia, and not confined to those cells that will undergo apoptosis. As genotoxic stresses engage a range of cell signaling pathways, it was possible that the enhanced IL-8 production that we observed in the neighbour cells reflected increased genotoxic stress in those cells, rather than a response to the retained apoptotic cells. In other words, E-cadherin KD might have enhanced the genotoxic stress of etoposide more generally, as well as inhibiting extrusion of apoptotic cells.

To test this possibility, we evaluated the DNA damage induced by etoposide by staining for gH2X, a classic marker of double-stranded DNA breaks. Both control and E-cadherin KD cells showed a significant increase in fluorescent intensity of nuclear gH2X staining after 24 hrs etoposide treatment (Fig S3B, C), and this was evident in most of the nuclei, supporting the idea that genotoxic damage was global. However, the increase in gH2X was similar in E-cadherin KD cells compared with controls (Fig S3B, C), suggesting that E-cadherin KD did not sensitize cells to DNA damage by etoposide. Consistent with this, we did not detect a significant difference in the proportion of cells that underwent apoptosis in time-lapse movies of control and E-cadherin KD cultures (Fig S3D). Nor did the production of IL-8 correlate with regional differences in DNA damage. We tested this by co-staining etoposide-treated cells for both IL-8 and gH2X, comparing gH2X intensity in cells that were IL-8 positive or IL-8 negative. There was no difference in DNA damage between these two groups (Fig S3E, F). Overall, we conclude that the enhanced IL-8 production observed in neighbour cells was not due to direct genotoxic stress in those cells. Instead, we considered the possibility that retained apoptotic cells might signal to stimulate IL-8 production by their neighbours.

### ATP signals from apoptotic cells to neighbour cells

How could retained apoptotic cells stimulate IL-8 production in their neighbours? One possibility was mechanical communication, as apoptotic cells become hypercontractile and stimulate tension-sensitive mechanotransduction pathways in their neighbours^26^. We tested this by asking if mimicking hypercontractility could stimulate IL-8 production regardless of other features of apoptosis. To do this, we transiently transfected E-cadherin KD monolayers with constitutively active ROCK (CA-ROCK), which induces cell contractility independently of apoptosis^25,26,38,39^, using an optimised transfection protocol that achieved 5-10% transfection efficiency to create sporadic hypercontractile cells within the confluent monolayer. However, CA-ROCK did not stimulate IL-8 production in E-cadherin KD monolayers (Fig 4A). This implied that a chemical signal from apoptotic cells might be required to stimulate IL-8 production in their neighbours.

**Fig 4.**
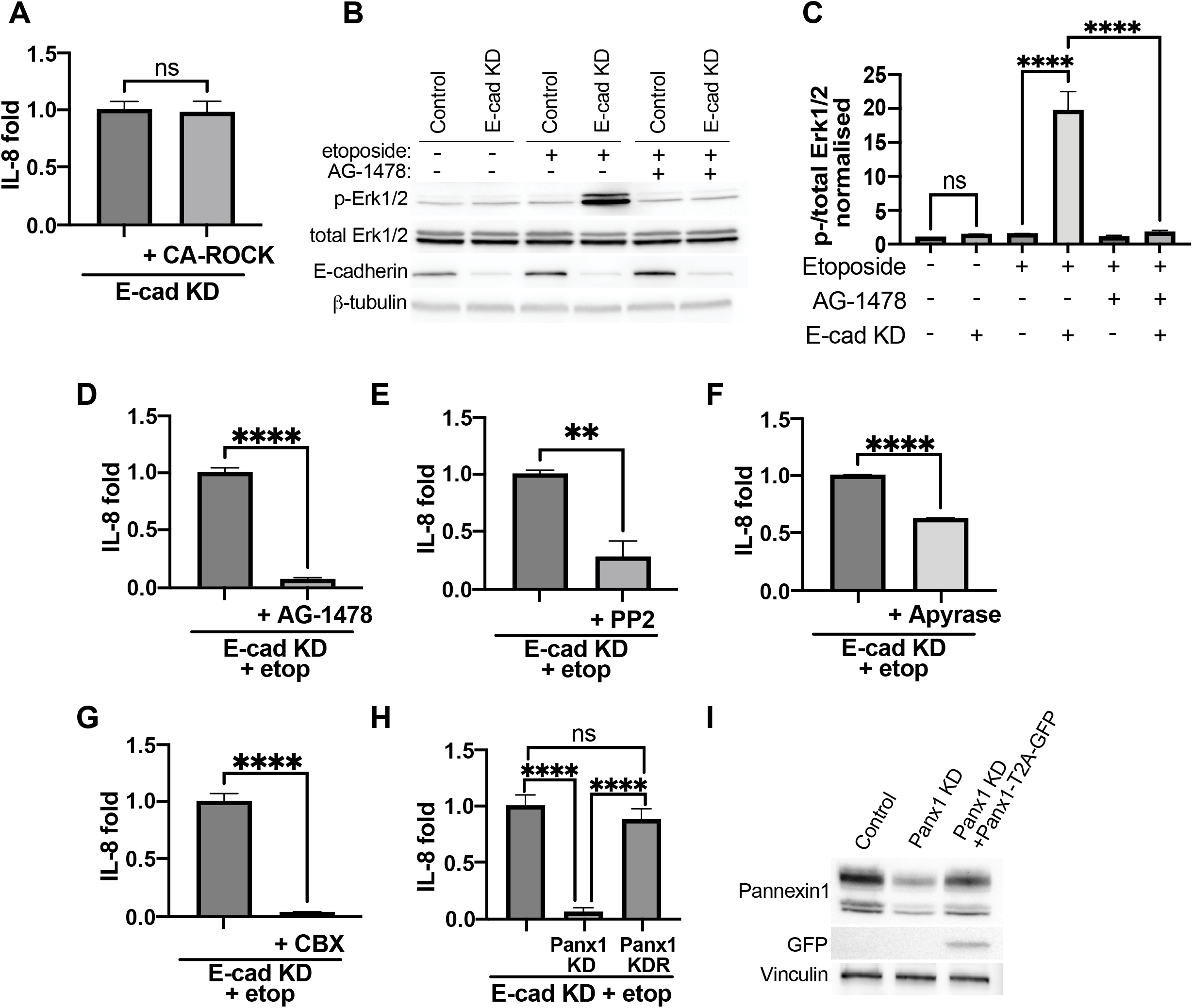
ATP derived from retained apoptotic cells is essential for IL-8 production. **A**. Fold IL-8 change upon mosaic transfection of E-cadherin knock-down (E-cad KD) cells with constitutively active ROCK (CA-ROCK). Values are normalised to E-cad KD. **B-C**. Immunoblotting (B) and quantification (C) of phospho- and total-Erk1/2 in control and E-cadherin KD cells treated with 250μM etoposide and EGFR inhibitor AG-1478 (1μM, 12hrs). **D-G**. Change in IL-8 production (fold, normalised to E-cadherin KD) upon treatment with (D) AG-1478 (1μM, 24hrs); (E) PP2 (10μM, 24hrs); (F) Apyrase (10U/ml, 24hrs); (G) carbenoxolone (CBX, 50μM, 24hrs). Experiments were performed on E-cadherin knockdown background, cells were treated with etoposide (250μM, 24hrs). **H-I**. Impact of pannexin 1 knock-down and its re-expression on IL-8 level (H) released from E-cadherin knock-down cells treated with etoposide (250μM, 24hrs). (I) - representative immunoblots of samples used for quantification in (H). All data are means ± SEM; ns, not significant, **p < 0.01, ***p < 0.001, ****p < 0.0001; calculated from n≥3 independent experiments analysed with Student’s t-test (A, D-G) or one-way ANOVA Dunnett’s multiple comparisons test (C, H).

Epidermal growth factor receptor (EGFR) tyrosine kinase signaling is often a key element that activates NF-κB in epithelial cells in response to extracellular stimuli^40–44^. This appeared to be engaged when apoptosis was induced in E-cadherin KD monolayers, as pERK levels increased and were blocked by the selective EGFR inhibitor Tyrphostin AG-1478 (Fig 4B, C). Furthermore, stimulation of IL-8 in these assays was also inhibited by AG-1478 (Fig 4D). We reasoned that candidate molecules which might be released from apoptotic cells to stimulate IL-8 production in their neighbours would engage the EGFR-NF-κB pathway.

Of potential candidates, ATP is released from apoptotic cells^45^ and can stimulate the Src-dependent transactivation of EGFR^46^ to induce IL-8 synthesis. A role for this mechanism in our system was supported by the observation that the Src inhibitor PP2 blocked IL-8 production when E-cadherin KD cells were stimulated with etoposide (Fig 4E). To further test a role for ATP, we treated E-cadherin KD cultures with apyrase to degrade extracellular ATP. Apyrase reduced the IL-8 response to etoposide, supporting a role for extracellular ATP (Fig 4F). As ATP is released from apoptotic cells through Pannexin 1 channels, we then treated cultures with carbenoxolone (CBX) to block Pannexin 1 channels. This significantly dampened the IL-8 response to etoposide in E-cadherin KD monolayers (Fig 4G). As CBX can also block other channels, such as connexins and hemiconnexins, we used RNAi to test a specific requirement for Pannexin 1. Pannexin1 siRNA robustly blocked etoposide-stimulated IL-8 production in E-cadherin KD cells, and this was restored when we coexpressed an RNAi-resistant Pannexin 1 transgene (Fig 4H,I). Together, these identify the release of ATP as playing a major role to induce IL-8 production in neighbours of apoptotic cells. They suggested that epithelia have an intrinsic pro-inflammatory cell-cell signaling circuit, where ATP released through Pannexin 1 channels from apoptotic cells induced IL-8 production in neighbour cells.

To reinforce this model, we then targeted the ATP receptor responsible for stimulating IL-8 production. We focused on P2Y2 and P2Y11, the purinergic receptors for extracellular ATP, and depleted each of these receptors by siRNA in E-cadherin KD monolayers. While P2Y2 KD had no effect on production of IL-8, the IL-8 response to etoposide was strongly suppressed by P2Y11 KD (Fig 5A). This was confirmed with the selective P2Y11 antagonist NF-157, which also inhibited IL-8 production (Fig 5B). Therefore, P2Y11 was principally responsible for transducing ATP signaling to stimulate IL-8. Interestingly, P2Y11 is reported to localize selectively to the basolateral surface of Caco-2 cells^47^. We confirmed this by expressing a P2Y11-GFP transgene in Caco-2 cells whose endogenous ZO-1 was tagged with mCherry by CRISPR/Cas9 gene editing. P2Y11-GFP localized to basolateral surfaces below the ZO-1-marked tight junctions (Fig 5C). Furthermore, apically-extruded apoptotic cells in control cultures were separated from the basolateral P2Y11-GFP by tight junctions. However, in E-cadherin KD monolayers the retained apoptotic cells remained directly opposed to the neighbour cell membranes bearing P2Y11-GFP (Fig 5D). Therefore, apical extrusion had the capacity to segregate apoptotic cells secreting ATP from their receptors, and this was lost when apical extrusion failed.

**Fig 5.**
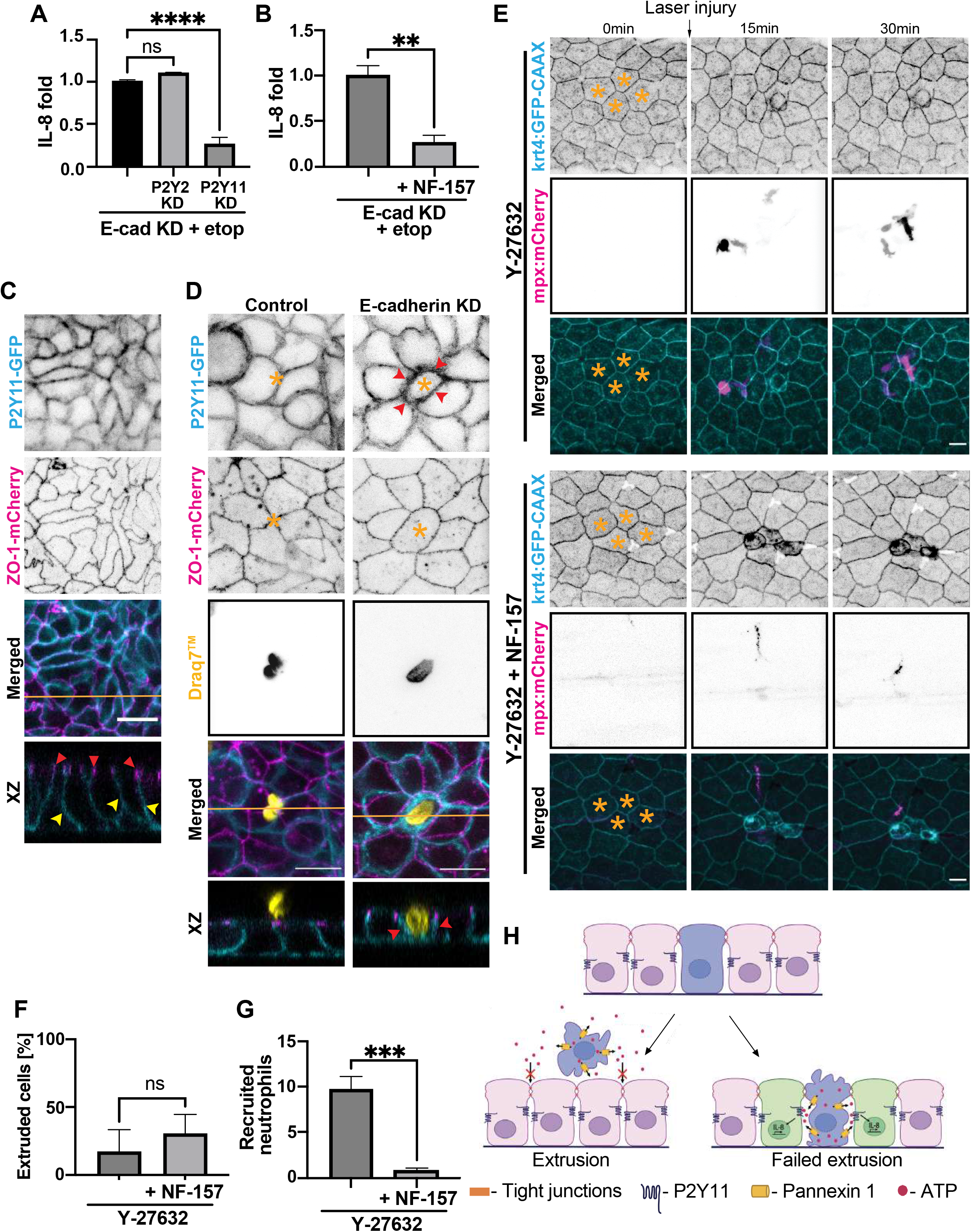
ATP-P2Y11 signalling axis stimulates IL-8 production upon failure of apoptotic extrusion. **A**. Impact of P2Y2 and P2Y11 knock-down on IL-8 level released from E-cadherin knockdown cells treated with etoposide (250μM, 24hrs). **B**. Impact of P2Y11 inhibitor NF-157 (5μM, 24hrs) on IL-8 level released from E-cadherin knock-down cells treated with etoposide (250μM, 24hrs). **C**. Representative images showing localisation of P2Y11-GFP transgene relative to endogenous mCherry-tagged ZO-1. P2Y11-GFP (yellow arrowheads) localises underneath ZO-1-mCherry (red arrowheads). Yellow line - location of the XZ view. **D**. Representative images showing localisation of P2Y11-GFP transgene upon successful (Control) and failed (E-cadherin KD) extrusion. Red arrowheads – accumulation of P2Y11 receptor around the retained apoptotic cell; yellow asterisk – apoptotic cells; yellow line - location of the XZ views. **E**. Selected frames form live-imaging of peridermal apoptosis and neutrophil recruitment to the sites of apoptosis in 4dpf zebrafish periderm. Prior to imaging zebrafish were treated with 45μM Y-27632 and 5μM NF-157 (16hrs). Yellow asterisk – apoptotic cells. **F**. Percentage of extruded apoptotic cells upon 45μM Y-27632 and 5μM NF-175 treatment (16hrs respectively) in 4dpf zebrafish. **G**. Number of neutrophils recruited to the sites of apoptosis within 30 min following laser microinjury of zebrafish peridermal cells. Experiments were performed on 4dpf zebrafish. Prior to imaging zebrafish were treated with 45μM Y-27632 and 5μM NF-175 (16hrs respectively). **H**. Schematic diagram: ATP derived from extruded apoptotic cells cannot access P2Y11 receptor located underneath tight junctions; upon failure of apoptotic extrusion ATP derived from retained apoptotic cells is able to access and activate P2Y11 receptor on surrounding healthy epithelial cells, stimulating them to produce IL-8. Scale bars: 15μm. XY panels are maximum projection views from middle z-stacks (P2Y11-GFP C, D) or all z-stacks (rest of images). All data are means ± SEM; ns, not significant, **p < 0.01, ***p < 0.001, ****p < 0.0001; calculated from n≥3 independent experiments analysed with one-way ANOVA Dunnett’s multiple comparisons test (A) or Student’s t-test (B, F, G).

Finally, we used NF-157 as a tool to confirm that ATP signaling drives the inflammatory response to retained apoptotic cells in zebrafish embryos. The double-positive offspring of *Tg(krt4:GFP-CAAX*)^uqay1^ and *Tg(mpx:mCherry*)^uqay3^ lines were treated with Y-27632 to disable apical extrusion with, or without, NF-157 to block ATP signaling before laser micro-irradiation assays. Apoptotic cells were retained when extrusion was blocked irrespective of whether ATP signaling was also inhibited (Fig 5F). However, addition of NF-157 significantly reduced the number of neutrophils that were recruited to the retained apoptotic cells (Fig 5E,G, Video S2). Therefore, ATP signaling is critical for epithelia to activate acute inflammation in response to apoptosis when apical extrusion fails.

## Discussion

Many of the anti-inflammatory responses to apoptosis in homeostasis have been elucidated from the study of cells, such as lymphocytes, that exist as single, isolated cells protected from their external environment by epithelial barriers. Our current findings reveal that, contrary to this traditional model, apoptosis in epithelia provokes acute inflammation through a intrinsic program that reflects the inherently communal nature of these tissues and their role as biological barriers in the body. Thus, an acute inflammatory response was provoked when apical extrusion was disabled and apoptotic cells retained in the epithelium. Evidently, neither the cell-intrinsic process of apoptotic fragmentation nor alternative homeostatic mechanisms, such as apoptotic clearance, could compensate to prevent inflammation when extrusion failed. Our findings indicate that this is because epithelia are hard-wired to elicit inflammation in response to apoptosis. This arises from a juxta-apoptotic signaling pathway, where ATP secreted from retained apoptotic cells stimulated the synthesis and secretion of IL-8 by their neighbours, leading to the recruitment of neutrophils (Fig 5H). Therefore, in contrast to the many active anti-inflammatory mechanisms that are known to be elicited by apoptosis, epithelia are wired to engage a pro-inflammatory response.

Membrane breakdown with release of internal cellular contents can directly engage immune responses^2^, and this direct mechanism has been invoked to explain how apoptotic cells may directly provoke inflammation if they persist^48,49^. In contrast, our data identify the neighbours of retained apoptotic cells as critically responsible for eliciting acute inflammation. Thus, IL-8 signaling was responsible for recruiting neutrophils in response to apoptosis when extrusion was blocked in zebrafish embryos and cell culture. Strikingly, upregulation of IL-8 synthesis did not occur throughout the whole monolayers, but was confined to the near-neighbours of the retained apoptotic cells, indicating that it was a local community response to the apoptotic cells. Importantly, juxta-apoptotic upregulation of IL-8 was abrogated when we targeted extracellular ATP, its P2Y11 receptor, or the pannexin channels that mediate its release from apoptotic cells. ATP has long been recognized as a canonical “find-me” signal whose release by apoptotic cells directly recruits immune cells^50^. Our data indicate that in epithelia it also works indirectly, to mediate the juxta-apoptotic upregulation of IL-8 by the epithelium itself.

Channels such as pannexins would be expected to release ATP from apoptotic cells irrespective of whether or not they were retained in the monolayer. How, then, can apical extrusion prevent the juxta-apoptotic activation of IL-8 secretion, especially as extruded apoptotic cells often persist at the apical surfaces of the epithelium? We suggest that this can be explained by the basolateral localization of the P2Y11 receptors responsible for stimulating IL-8 synthesis. ATP released from extruded apoptotic cells, with a 14Å effective diffusional diameter^51^, would be prevented from accessing P2Y11 receptors by the tight junctions (pore size <8Å^52,53^) that seal the epithelium as the apoptotic cell is being expelled. However, when apical extrusion was disrupted, the retained apoptotic cells remained contiguous with the P2Y11-enriched basolateral surfaces of their immediate neighbours, which would allow secreted ATP to directly access its receptors to stimulate IL-8 synthesis. This proximity may also explain why IL-8 is upregulated as a local response in the epithelium. As well, tight junctions often disassemble when apoptotic extrusion is compromised^26^, which would encourage leakage of ATP from apoptotic cells out of the epithelium, further limiting the distribution of ATP to the immediate locale.

Why would epithelia have developed a pro-inflammatory response to the presence of apoptotic cells, in contrast to the anti-inflammatory strategies evinced by other responders, such as macrophages^20–22^? One possibility is that this mechanism represents a fail-safe, to ensure that apoptotic cells unable to be extruded from the monolayer are targeted by neutrophils that orchestrate an efferocytotic response to clear the retained apoptotic cells. Although neutrophils are not well-characterized as phagocytes, they can recruit macrophages as professional mediators of apoptotic clearance^54^. However, in zebrafish embryos, neutrophils were recruited within minutes of inducing apoptosis, suggesting that the para-apoptotic inflammatory response is an active early-immediate response of some kind.

Another possibility lies in the well characterised anti-microbial effector functions of neutrophils. Extrusion failure causes local defects in the epithelial barrier that would constitute sites for microbial ingress, because retained apoptotic cells contract and physically detach from their neighbours^26^. Thus, the para-apoptotic IL-8 response may be an early-warning mechanism that helps identify nascent barrier defects, recruiting neutrophils as a defence response to barrier failure and the possibility of microbial breach. In other words, if the epithelial barrier is the first line that defends the body from its external environment, the neutrophil recruitment that we observed would constitute a mechanism for the failing barrier to engage a second line of defence. However, having a pro-inflammatory circuit hard-wired into epithelia carries the risk that sporadic apoptosis, which occurs frequently in daily life^11^, would provoke excessive inflammation. We propose that apical extrusion mitigates this risk and prevents premature inflammation by physically eliminating apoptotic cells from the body before they can provoke IL-8 signaling in their neighbours. Of note, apical extrusion can be compromised by diverse factors, including changes in tissue mechanics^55,56^ and metabolism^57^. Therefore, extrusion failure that prevents apoptotic cells from being eliminated may provide a new mechanism to increase the risk of inflammation in epithelial tissues.

## Acknowledgements

We thank our colleagues for their support and advice throughout this project. Our work was supported by grants from the National Health and Medical Research Council of Australia (1136592 and 1163462 to A.S.Y; 2009075 to K.S.; APP1194406 to M.J.S.) and Australian Research Council (DP220103951 to A.S.Y). Microscopy was performed at the ACRF/IMB Cancer Research Imaging Facility created with the generous support of the Australian Cancer Research Foundation.

## Author contributions

Conceptualization, K.D. and A.S.Y.; investigation, K.D., J.B.P., D.R., D.C.R., S.V., F.L.; resources, M.J.S, K.S.; funding acquisition, A.S.Y.; writing, K.D., K.S. and A.S.Y.; supervision, A.S.Y.

